# Tissue specific muscle extracellular matrix hydrogel improves skeletal muscle regeneration *in vivo* over non-matched tissue source

**DOI:** 10.1101/2020.06.30.181164

**Authors:** Jessica L. Ungerleider, Monika Dzieciatkowska, Kirk C. Hansen, Karen L. Christman

**Affiliations:** Department of Bioengineering, Sanford Consortium for Regenerative Medicine, University of California, San Diego, La Jolla, CA, USA; Department of Biochemistry and Molecular Genetics, School of Medicine, University of Colorado, Aurora, CO, USA

**Author notes:** Corresponding, University of California San Diego, Department of Bioengineering, Sanford Consortium for Regenerative Medicine, 2880 Torrey Pines Scenic Drive, La Jolla, CA 92037. Author contributions: JLU – design study, wrote paper, conduct experiments and analyze and interpret data; KLC – obtained funding, design study, interpret data, wrote paper. MK - conduct experiments and analyze and interpret data. KH interpret data, edited paper.

**Keywords:** Extracellular matrix, hydrogel, skeletal muscle, regeneration

## Abstract

Decellularized extracellular matrix (ECM) hydrogels present a novel, clinical intervention for a myriad of regenerative medicine applications. The source of ECM is typically the same tissue to which the treatment is applied; however, the need for tissue specific ECM sources has not been rigorously studied. We hypothesized that tissue specific ECM would improve regeneration through preferentially stimulating physiologically relevant processes (e.g. progenitor cell proliferation and differentiation). One of two decellularized hydrogels (tissue specific skeletal muscle or non mesoderm-derived lung) or saline were injected intramuscularly two days after notexin injection in mice (n=7 per time point) and muscle was harvested at days 5 and 14 for histological and gene expression analysis. Both injectable hydrogels were decellularized using the same detergent and were controlled for donor characteristics (i.e. species, age). At day 5, the skeletal muscle ECM hydrogel significantly increased the density of Pax7+ satellite cells in the muscle. Gene expression analysis at day 5 showed that skeletal muscle ECM hydrogels increased expression of genes implicated in muscle contractility. By day 14, skeletal muscle ECM hydrogels improved muscle regeneration over saline and lung ECM hydrogels as shown through a shift in fiber cross sectional area distribution towards larger fibers. This data indicates a potential role for muscle-specific regenerative capacity of decellularized, injectable muscle hydrogels. Further transcriptomic analysis of whole muscle mRNA indicates the mechanism of tissue specific ECM-mediated tissue repair may be immune and metabolism pathway-driven. Taken together, this suggests there is benefit in using tissue specific ECM for regenerative medicine applications.

## 1. Introduction

Acute and chronic muscle injuries are common ailments, often caused by physical trauma or as a co-morbidity of lifestyle diseases. Skeletal muscle repair is a complex regenerative process, requiring exquisite spatiotemporal coordination of many cellular subtypes in a complex inflammatory milieu. Dysregulation of this process prevents functional repair of muscle due to improper myofibril function and organization. Activation of quiescent stem cell populations, rapid proliferation of myocytes, and differentiation into multi-cellular myotubes is required for this repair to occur. However improper myogenesis and inflammation is associated with aging and various disease states [1]. A critical need exists for therapeutic interventions to guide this process.

Extracellular matrix (ECM) hydrogels, most commonly formed through enzymatic digestion of decellularized ECM, show great promise in a variety of regenerative medicine applications [2-4]. Their retention of the complex distribution of native ECM proteins and proteoglycans is able to effectively guide cell fate decisions. Additionally, while ECM hydrogels do not retain the tissue’s native 3D ultrastructure, they can reassemble into a hydrogel at physiological temperature, pH, and salt concentration and form a nanofibrous matrix with similar architecture to native tissue [5]. Due to their ability to form a hydrogel supportive matrix *in situ*, injectable ECM hydrogels are translationally attractive for applications where minimally invasive delivery is desirable (injections over a wide area or difficult to access location). ECM hydrogels have shown efficacy in numerous preclinical models [2, 3] including myocardial infarction [6, 7], peripheral artery disease [8], and *in vitro* 3D microenvironmental drug screening models [9], and were recently translated into patients in a Phase I clinical trial in myocardial infarction patients [10].

A standard approach is to use an ECM hydrogel where the source tissue that is decellularized is typically the same tissue to which the treatment is applied. While studies have investigated the merit of using tissue specific ECM sources *in vitro* [11-17], the importance of tissue specificity has not been definitively assessed *in vivo.* Some groups have compared different ECM materials for esophageal [18] or skeletal muscle repair [8], but the control “non-matched” ECM sources were not properly controlled for form (intact patch versus solubilized hydrogel) or ECM age and decellularization method, thus complicating the question of young versus old ECM and effect of decellularization method on biological outcomes *in vivo*. We therefore sought to evaluate the importance of tissue specificity more rigorously by controlling for ECM species, age, and decellularization detergent. We hypothesized, in a muscle regeneration model, that a tissue specific ECM hydrogel would improve regeneration and function through preferentially stimulating physiologically relevant processes (e.g. progenitor cell proliferation and differentiation). In this study we show that skeletal muscle ECM hydrogels benefit skeletal muscle regeneration over a non-matched tissue source (lung) via stimulation of skeletal muscle progenitor recruitment and increased muscle growth.

## 2. Materials and Methods

### 2.1 ECM fabrication and characterization

ECM hydrogels were fabricated and characterized using previously published methods [8, 19]. Briefly, skeletal muscle or lung tissue from 3 month old pigs was harvested and chopped into small pieces. Tissue was rinsed in ultrapure water for 30 minutes followed by ∼4 days of rinsing in 1% (skeletal muscle) or 0.1% (lung) sodium dodecyl sulfate with periodic solution changes. Skeletal muscle material was finally rinsed in isopropanol for 18 hours to remove interstitial fat. Final material was lyophilized, milled into a fine powder, and digested for 48 hours in 1 mg/ml pepsin in 0.1M HCl. Digests were neutralized and lyophilized at 6 mg/ml. ECM targeted, QconCAT proteomics were performed on the milled material pre-digestion as previously described [20]. Rheometry was performed using an AR-G2 rheometer on 500 µL gels at 37 °C with a 1000 µm gap height and 2.5% strain for shear moduli and on 200 µL liquid ECM with a 500 µm gap height at 25 °C for complex viscosity measurements. DNA and sulfated glycosaminooglycan (sGAG) content were determined as previously described [19].

### 2.2 NTX injury

This work was carried out in accordance with the Institutional Animal Care and Use Committee at UC San Diego. Male 8 week old C57BL/6 animals were injured using a notexin acute muscle injury model. Notexin (10µg/mL, Latoxan) was injected in a single injection of 20 µL into the mid-belly of the right tibialis anterior (TA) muscle. Left TAs served as noninjected and uninjured controls. After 2 days, animals were injected with a single 20 µL injection of either saline, skeletal muscle ECM, or lung ECM (n=7 per group at each time point). Animals were then harvested either 3, 7, or 12 days post-treatment. TA muscles were laid on filter paper to orient fibers and flash frozen in liquid nitrogen. Muscles were then frozen again in OCT for cryosectioning along the short axis. During cryosectioning, tissue was either mounted on slides (10µm thick sections) or collected for RNA isolation at 3 locations spanning the muscle.

### 2.3 Immunohistochemistry

Cryosections were fixed with 4% paraformaldehyde and blocked with 1% bovine serum albumin, 0.3% Triton-X, and 5% goat serum. Primary antibodies against laminin (1:100, abcam), and Pax7 (1:2, DSHB) were incubated overnight at 4°C. Secondary antibodies (AlexaFluor goat IgG against rabbit [laminin], mouse [Pax7]) were incubated at 1:1000 for 1 hour at room temperature and nuclei were visualized with Hoechst 33342.

### 2.4 RNA sequencing

RNA was isolated using Trizol and cleaned using Qiagen on-column DNase clean-up per the manufacturer’s instructions. The Institute for Genomic Medicine at UCSD conducted all quality checks, library prep, and sequencing of the RNA samples. Briefly, RINe was measured using an Agilent Bioanalyzer, libraries were prepared using Illumina’s mRNA stranded library prep kit, and libraries were sequenced to a depth of approximately 25 million reads on a HiSeq flow lane. Quality of sequencing was verified with FastQC. Fastq files were aligned to the Ensembl mouse genome (Grcm38 version 84) using HiSat 2 [21] and counted using StringTie [22, 23]. Counts were statistically analyzed using the DESeq2 package [24]. Differentially expressed genes were defined as genes with adjusted Benjamini-Hochberg p-value less than 0.1 and absolute value of log fold change greater than 0.6. Differentially expressed genes were assigned to biological pathways using DAVID functional annotation clustering [25, 26].

## 3. Results

### 3.1 Tissue-of-origin alters composition of injectable ECM hydrogels

To understand how tissue-of-origin alters muscle repair, we developed injectable hydrogels from two orthogonal sources – skeletal muscle and lung. Lung was chosen as a non-matched tissue control as it is distinct from skeletal muscle in both developmental origin and tissue function. Both injectable hydrogels were decellularized using the same detergent (SDS) and were controlled for donor characteristics (i.e. species, age).

Common characterization assays – DNA content, sulfated glycosaminoglycan content, and shear rheology – show that all three ECM hydrogels were sufficiently decellularized and had similar viscoelastic properties (Figure 1). Fifty-three of the most common ECM and cellular contaminant proteins were quantified in the decellularized ECM using stable isotope labeled quantitative concatenated (QconCAT) peptides and targeted LC-MS analysis (Table 1). While both ECM scaffolds largely contained collagen I (65-80% by molar fraction), each scaffold had a complex and unique composition of ECM proteins. Importantly, collagen VI composition is different between the materials, which has been shown to have an effect on skeletal muscle progenitor cell renewal *in vivo* [27].

**Table 1:**
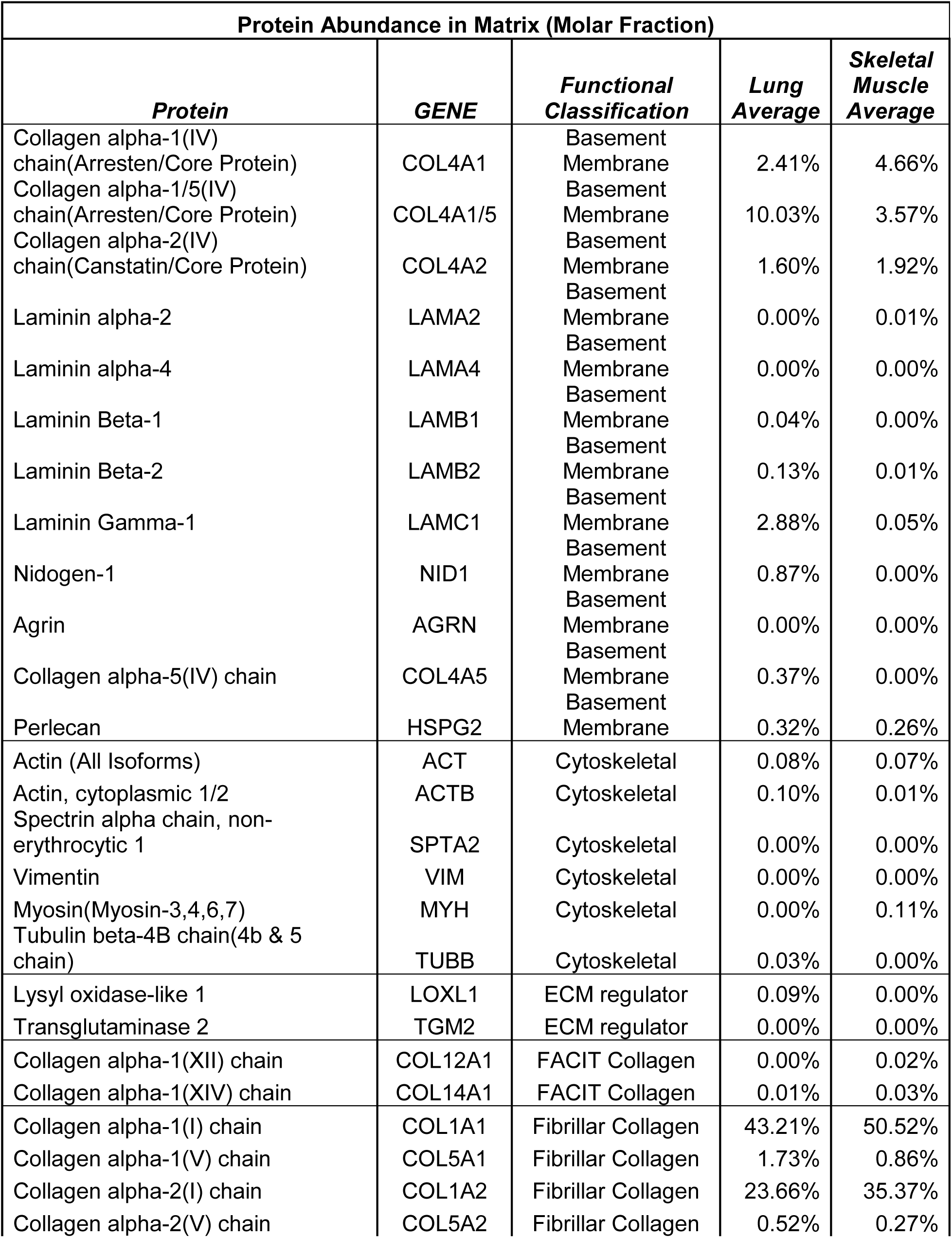
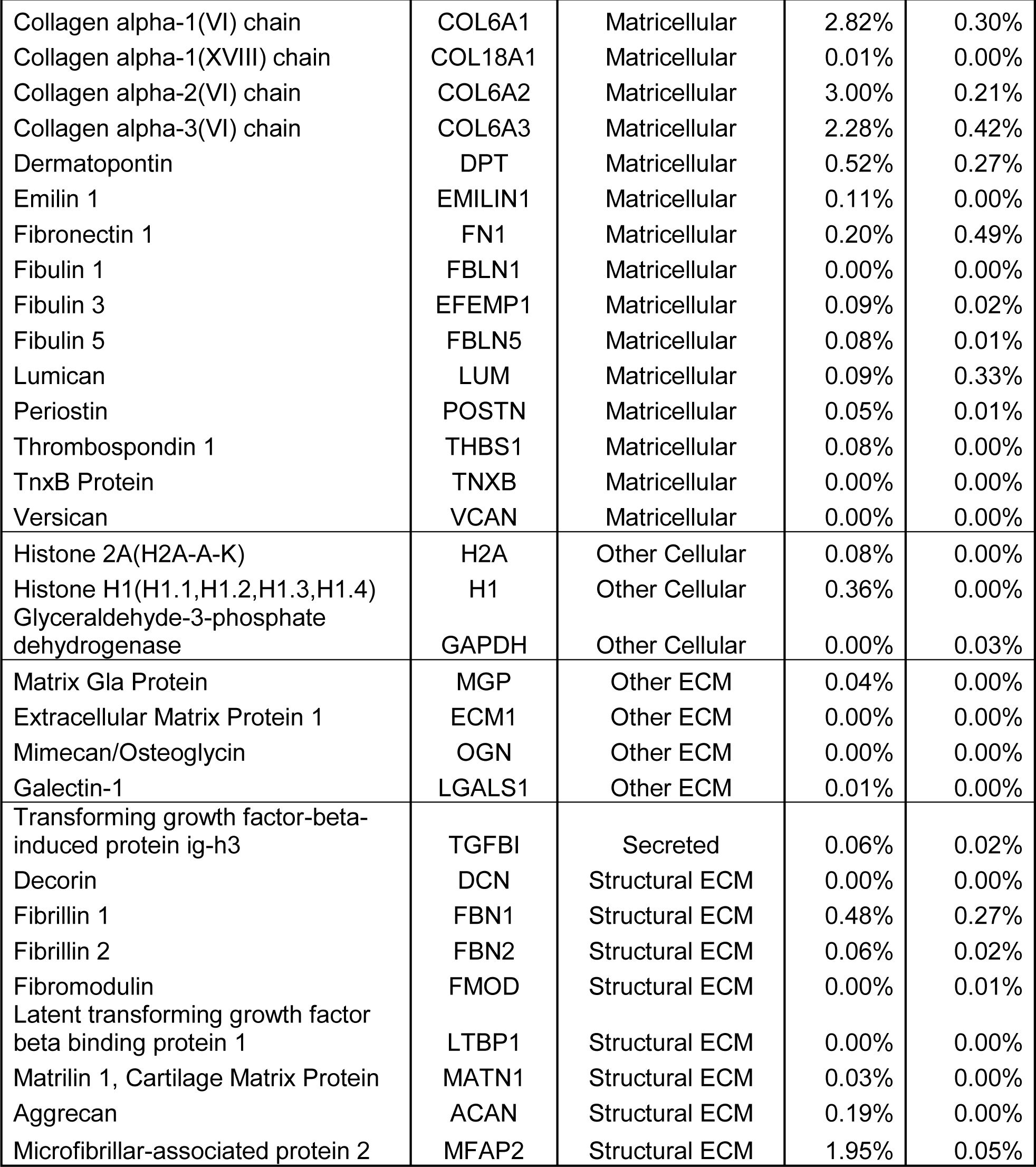
Quantitative proteomics of decellularized ECM from different tissues of origin.

**Figure 1.**
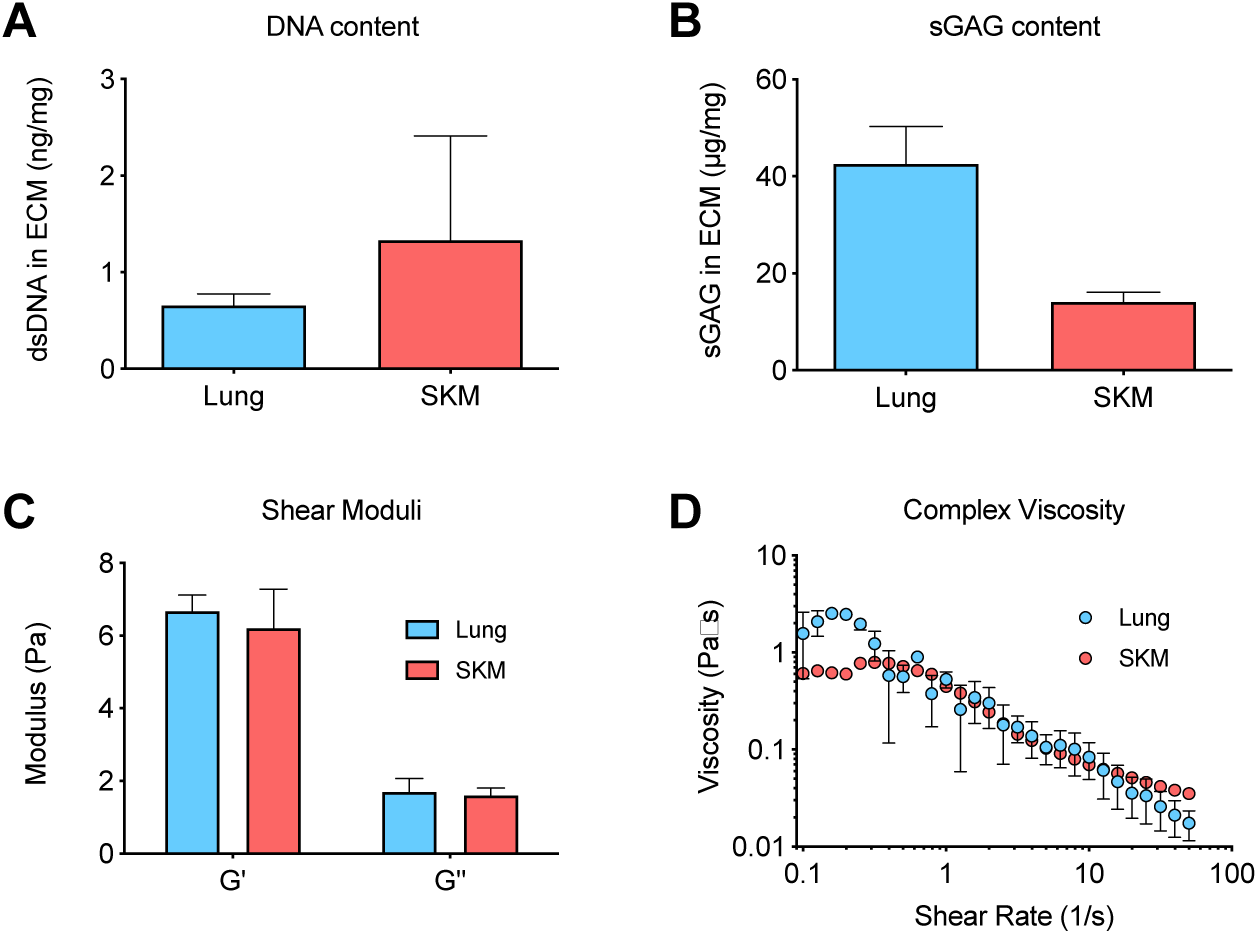
ECM hydrogel characterization. (A) DNA content remaining in decellularized hydrogels. (B) sulfated glycosaminoglycan (sGAG) content remaining in decellularized hydrogels. (C) Storage and loss moduli at 1 rad/s and 2.5% strain demonstrating gel formation. (D) Complex viscosity indicates materials are shear thinning.

### 3.2 Muscle-derived injectable hydrogel increases muscle repair after NTX injury

To investigate the difference in regenerative capacity of the two hydrogels *in vivo*, one of the two decellularized hydrogels, tissue specific skeletal muscle or non-mesoderm-derived lung, or saline were injected intramuscularly two days after notexin injection in male adult C57BL/6 mice (n=7 per time point) and muscle was harvested at days 3, 7, and 12 post-treatment for histological and gene expression analysis.

At day 12, the skeletal muscle ECM hydrogel led to an increase in fiber cross sectional area compared to the lung ECM hydrogel and saline (average fiber areas – skeletal muscle ECM: 695 ± 94 µm^2^; lung ECM: 545 ± 65 µm^2^; saline: 564 ± 64 µm^2^) This increase was evident through a significant decrease in fibers of smaller areas and a shift in distribution towards larger areas (Figure 2). This result suggested that skeletal muscle ECM hydrogels was able to improve muscle regeneration over saline and lung ECM hydrogels.

**Figure 2.**
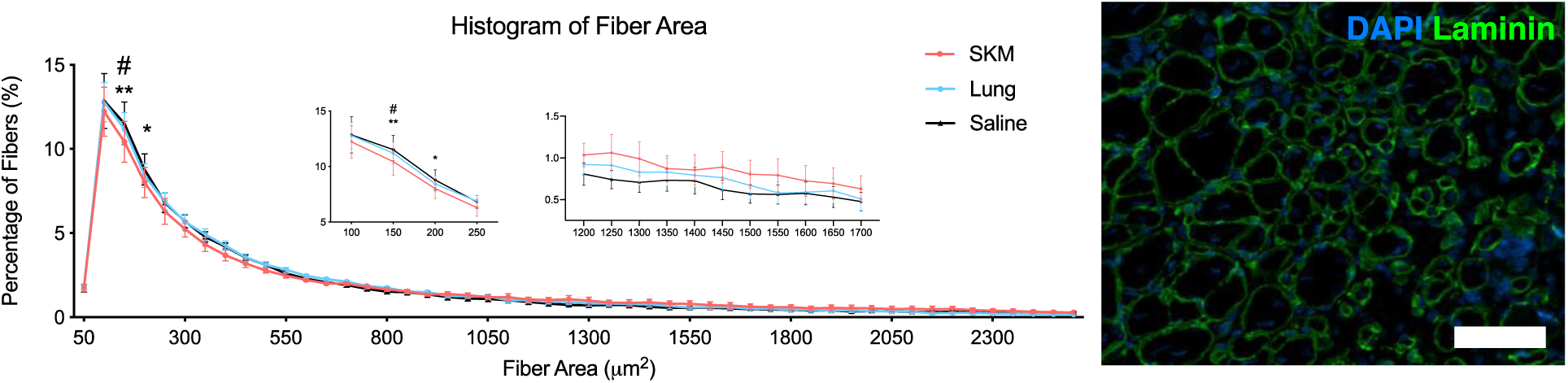
Skeletal muscle ECM hydrogel improves muscle fiber cross-sectional area over non-matched tissue source. A, Graph of fiber areas indicates the skeletal muscle ECM hydrogel (SKM) has fewer small fibers than saline and the lung ECM hydrogel (Lung). Insets show zoomed in sections of main graph with smaller fiber areas (left) and larger fiber areas (right). B, Representative image of fiber area quantified by laminin staining. *p<0.05 SKM versus saline, **p<0.01 SKM versus saline, #p<0.05 SKM versus Lung by two-way ANOVA with Fisher’s post-hoc test. Scale bar is 70 µm.

To better understand what processes could drive increases in muscle regeneration, we histologically-analyzed muscle repair processes during the regeneration process. At day 3, the skeletal muscle ECM hydrogel significantly increased the density of Pax7+ satellite cells in the muscle (Figure 3). This effect has also been previously shown in response to the skeletal muscle ECM hydrogel compared to collagen or saline [8, 28], but interestingly this effect was not seen with the non-matched lung ECM source. This further suggests that the skeletal muscle ECM composition is important for skeletal muscle progenitor activity *in vivo*.

**Figure 3.**
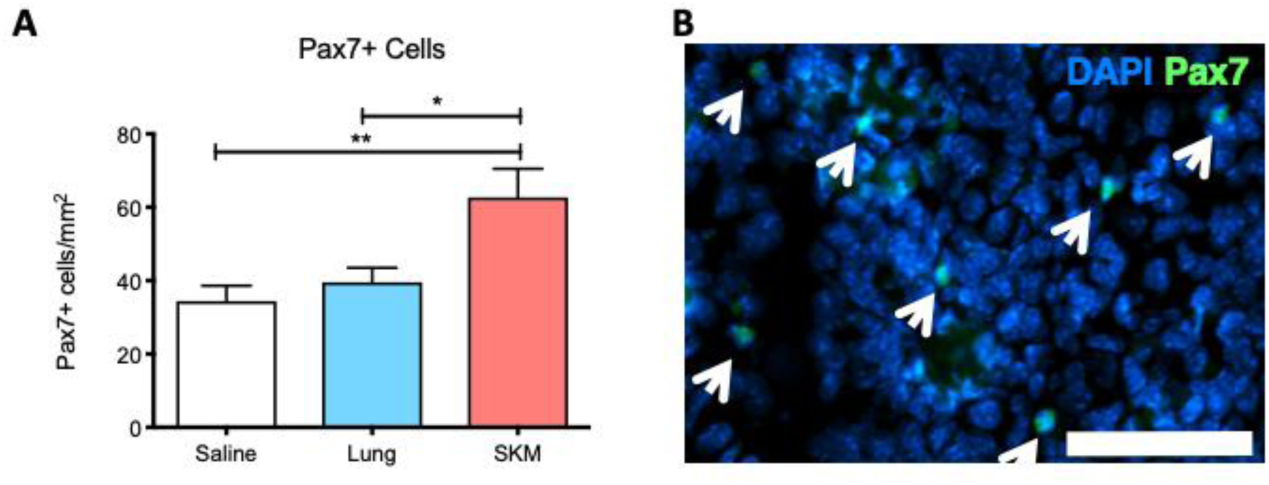
Skeletal muscle ECM hydrogel increases density of skeletal muscle progenitors. (A) Quantification of Pax7+ skeletal muscle progenitors. (B) Arrows indicate Pax7+ nuclei. *p<0.05, **p<0.01 using one-way ANOVA with Tukey’s post-hoc test. Scale bar is 70 µm. SKM = skeletal muscle ECM hydrogel. Lung = lung ECM hydrogel.

### 3.3 Gene expression indicates muscle hydrogel improves muscle contractility pathways

Differences in transcriptomic regulation due to material injection were investigated using RNAseq. Whole muscle RNA was isolated at 3 and 7 days post-injection, sequenced, and normalized to saline control injection. Differentially expressed (DE) genes relative to saline were biologically interpreted using DAVID functional annotation clustering analysis. Differentially regulated genes and pathways between skeletal muscle ECM hydrogel-saline and lung ECM hydrogel-saline were then compared to assess how the different biomaterials were altering the transcriptomic response *in vivo.* At day 3 post-injection, both materials clustered differently from saline using principal component analysis (PCA) (Figure 4). While there were 222 DE genes for skeletal muscle ECM hydrogel-saline and 552 DE genes for lung ECM hdyrogel-saline at day 3, this dropped down to 4 and 8 DE genes at day 7, respectively. Groups also did not cluster separately on PCA at day 7, suggesting most of the transcriptomic effects of ECM injection occurred during the first week post-injection. It has also been shown in previous transcriptomic studies that major differences due to ECM injection in skeletal muscle occur at shorter time points post-injection [8].

**Figure 4.**
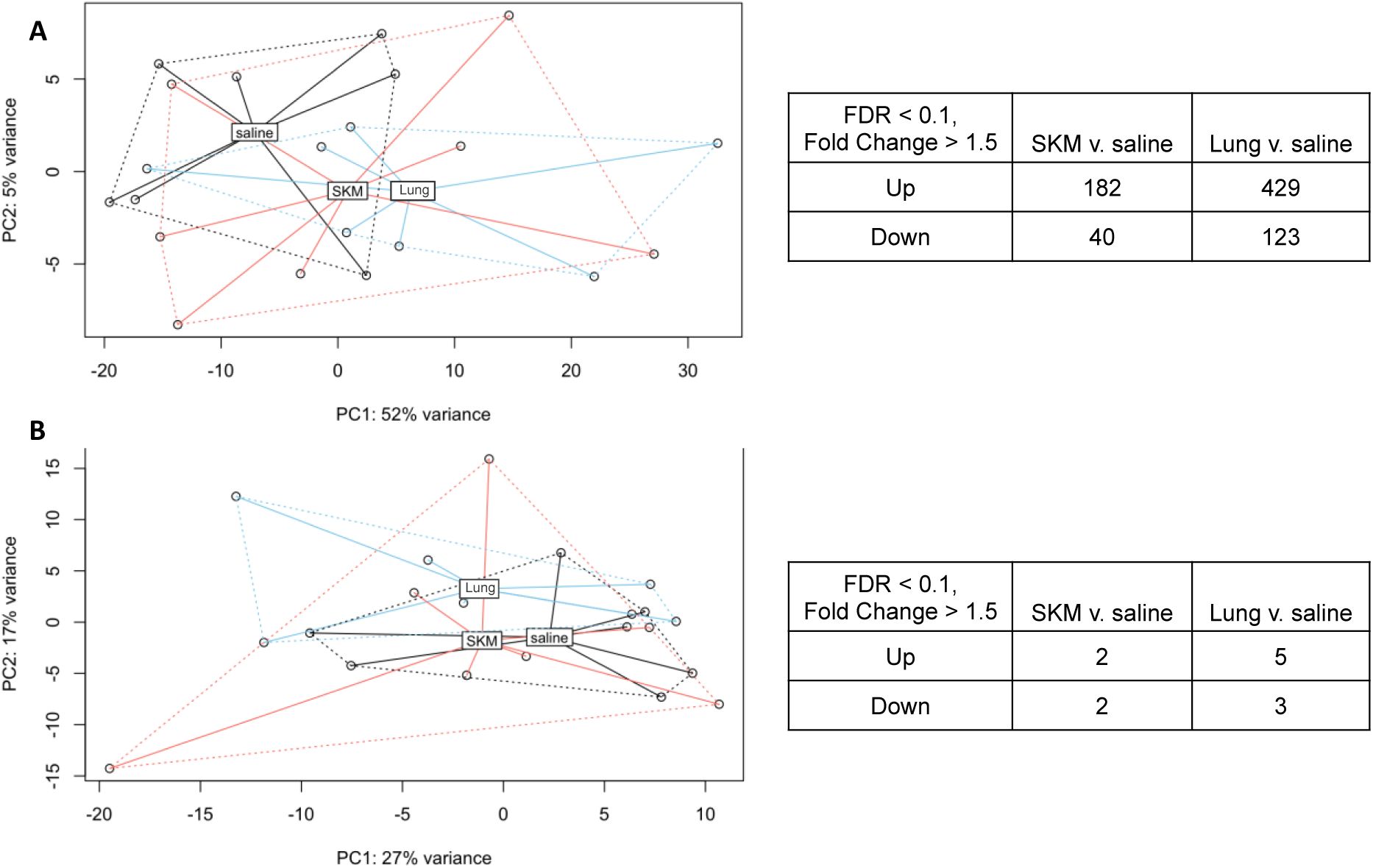
Differential expression analysis of notexin-injured and ECM hydrogel-treated muscles. Gene expression analysis was conducted using DESeq2 negative binomial generalized linear models. SKM = skeletal muscle ECM hydrogel. Lung = lung ECM hydrogel.

DAVID functional annotation clustering at day 3 post-injection shows that most DE pathways are related to metabolism, immune response, and skeletal muscle function (Table 2). Closer investigation of the overlapping and distinct DE genes was performed using Venn diagrams (Figure 5). Interestingly, the skeletal muscle ECM hydrogel showed, distinct from the lung ECM hydrogel, upregulation of genes related to skeletal muscle contraction (*Tnnt3, Tcap, Jsrp1, Mylk2*). This suggests a tissue specific effect of the muscle hydrogel on muscle contractility pathways, which is evidence that tissue specific ECM source is an important consideration in regenerative applications. Interestingly, the lung ECM hydrogel showed most distinct expression of genes related to metabolism and immune response, specifically upregulation of oxidative phosphorylation genes (ATP synthases *Atp5f1* and *Atp5j*, NADH dehydrogenases, and cytochrome c oxidases) and downregulation of inflammatory response genes (*Cd163, Adam8, S1pr3, Csf1, Il4ra*) relative to saline. Interestingly, some of these genes are implicated in the pro-regenerative response [29-32], suggesting the response to the lung ECM hydrogel may be more shifted to pro-inflammatory response at this time point. Overlapping pathways between the two materials include upregulation of c-type lectin (*Klr* family) and immune response (*Ccl5, Cd27*). Both gene sets are suggestive of an innate inflammatory response, which is expected at this early time point after injection of naturally derived ECM biomaterials before the shift towards a pro-remodeling phenotype [33, 34]. Interestingly, some signal/secreted transcripts were downregulated in both ECM groups (*Mmp9, Igtb3, Cxcl10, Osm*), which suggests regulation of ECM deposition and inflammatory response. However, as these genes are involved with many pathways, it is important to further validate their role in the tissue response to ECM hydrogels, especially since harnessing the inflammatory response is vital for successful skeletal muscle regeneration [31, 35, 36].

**Table 2:** DAVID functional annotation analysis of differentially expressed genes at day 3 post-injection. Top six up- and down-regulated pathways were reported by enrichment score.

| Skeletal muscle ECM hydrogel |  | Lung ECM hydrogel |  |
| --- | --- | --- | --- |
| Up-regulated vs. saline |  |  |  |
| Annotation Cluster | Enrichment Score | Annotation Cluster | Enrichment Score |
| C-type lectin, cell adhesion | 3.76 | C-type lectin, cell adhesion | 5.90 |
| Skeletal muscle contraction | 3.35 | Mitochondrion | 5.81 |
| Transmembrane | 2.49 | Oxidative phosphorylation | 4.80 |
| Disulfide bond/glycoprotein | 1.52 | Disulfide bond/signal/secreted | 3.51 |
| Ion channel activity | 1.37 | T cell receptor complex | 3.00 |
| Glycogen/carbohydrate metabolism | 1.20 | Transmembrane | 1.80 |
| Down-regulated vs. saline |  |  |  |
| Disulfide bond/signal peptide | 3.03 | Cadherin/cell adhesion | 6.85 |
| Cholesterol metabolic process | 1.88 | Signal/secreted/glycoprotein | 3.67 |
| Oxidation-reduction | 1.54 | Lipid metabolic process | 2.35 |
| Cytokine activity | 1.35 | Cytokine activity | 1.68 |
| Transmembrane | 1.28 | Oxidation-reduction | 1.65 |
| Protease binding | 1.20 | Cell division | 1.54 |

**Figure 5:**
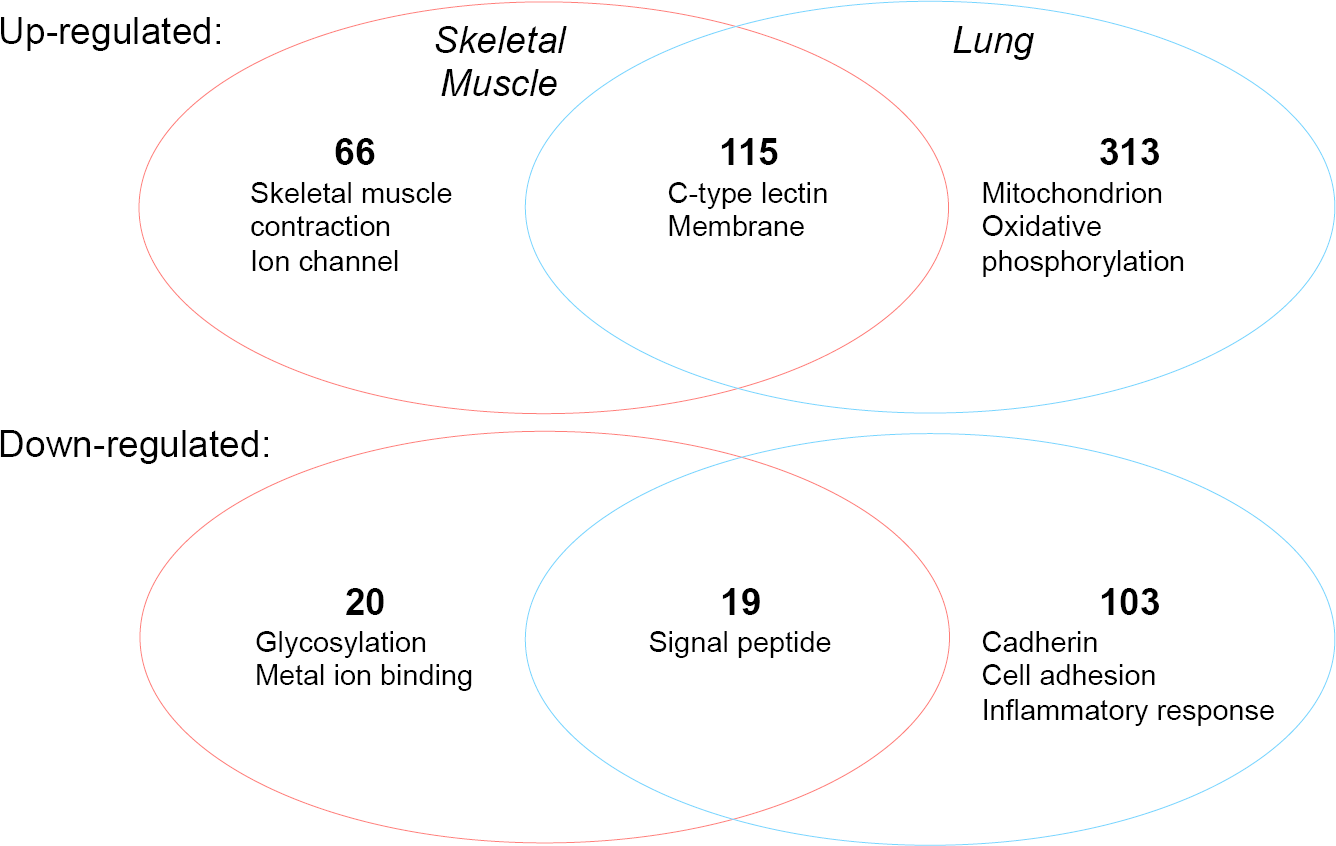
Overlapping and distinct differentially expressed genes between skeletal muscle and lung ECM hydrogels. Number of DE genes intersecting between SKM-saline and LNG-saline contrasts are shown as well as the top pathways. Pathways are gene ontology or Kegg pathways with the lowest p values as determined by DAVID at day 3 post-injection.

Overall, RNAseq analysis suggests that both ECM hydrogels induce non-specific effects such as differential regulation of metabolism and inflammatory response. However, there are some subtle differences in inflammatory cell response (i.e., downregulation of pro-regenerative genes in Lung), which may indicate a tissue-specific response in not only muscle regeneration and contractility response but also inflammation, which is known to have a lot of cross-talk with muscle regeneration pathways *in vivo.*[37]

## 4. Discussion

Decellularized tissues have long been used as scaffolds for regenerating or repairing damaged tissues. The prevailing thought is that by mimicking the endogenous ECM, therapeutic scaffolds can serve as not only a mechanical support but also as a biomimetic cue reminiscent of the healthy ECM. At first, these scaffolds were used generally as ECM mimetic scaffolds without thought for the specific tissue to which they were being applied. Urinary bladder matrix (UBM) and small intestinal submucosal (SIS) matrix were first applied as patches for encouraging tissue regeneration and improved healing in hernia repair, deep tissue burns, and wound dressings [38, 39]. They provided a rigid structural patch and showed bioactivity through integration of the patch with host tissue. Since then, many groups have adapted ECM hydrogels for regenerative medicine applications, using the ECM from the same tissue type which they are aiming to regenerate [3]. The importance of tissue specificity has been explored *in vitro* through comparison between single ECM components [13, 40] and ECM combinatorial screening approaches [41]. Lately, groups have been comparing ECM derived from different decellularized organs and have found that there are important tissue specific effects *in vitro* [9, 14, 16, 17]. Specifically, in the muscle, it has been shown that a tissue specific ECM improves differentiation and proliferation *in vitro* [40, 42]. Mechanistic studies where groups investigated which ECM components specifically were having an effect found that laminin and fibronectin were vital for myoblast migration and proliferation [43]. While these studies have importantly demonstrated that ECM is not interchangeable, this phenomenon has not been rigorously explored *in vivo*.

In this study, we sought to investigate the effect of tissue-specific decellularized ECM hydrogels on skeletal muscle regeneration. We first characterized the ECM hydrogels derived from adult porcine tissue sources, either tissue-specific skeletal muscle or functionally and developmentally non-specific lung. Both ECM hydrogels had similar mechanical properties, but biochemically they had distinct proteomic compositions. We next sought to determine the importance of tissue-specificity of decellularized hydrogels for *in vivo* regeneration in an acute muscle injury model, the notexin injury model, which is commonly used in the muscle regeneration field [44-46]. Muscle fiber cross-sectional area, an important metric of functional regeneration, was increased in the skeletal muscle ECM hydrogel treated animals at day 12 post-treatment. We also quantified Pax7+ satellite cells at day 3 post-injection, a timepoint of maximal satellite cell activation [45], and found that the skeletal muscle ECM hydrogel increased density of this cell type in the muscle over the lung ECM hydrogel and saline injected muscles. Finally, whole muscle transcriptomics revealed that most differences due to ECM injection (normalized to saline-injected control muscles) were seen at day 3 post-injection with upregulation of skeletal muscle contraction genes in the skeletal muscle ECM group and not in the lung ECM group.

The importance of tissue-specific ECM hydrogels on regenerative cell types and pathways has previously been rigorously studied *in vitro* [9, 13, 17, 42], but with limited confirmatory evidence *in vivo* [8, 18]. This study for the first time demonstrated the importance of tissue specificity *in vivo* while rigorously controlling for decellularization species, age, detergent, and mechanical properties of the resultant material. This is an important finding, because many non-specific decellularized ECM patches are routinely used for regenerative medicine such as SIS and UBM. These materials have shown efficacy *in vivo*, but our study suggests that tissue-specific materials can be more beneficial for regenerative medicine applications than non-organic specific ECM. Benefits seen with SIS and UBM *in vivo* could be explained by the non-specific pathways we consistently see being differentially regulated by ECM materials (i.e. inflammation and metabolism). The impact of these non-specific pathways should be further elucidated with detailed mechanistic studies. Depending on the application, the polarization of these non-specific pathways may be sufficient alone (e.g., in wound healing [47, 48]).

In this study both ECM hydrogels had similar mechanical properties so *in vivo* responses were likely due to biochemical effects induced by differences in hydrogel proteomic composition. ECM has been shown to be an important regulator of skeletal muscle satellite cell response *in vivo* [49]. For example, collagen VI is a basement membrane protein known to be important for the muscle satellite cell self-renewal [27]. However, this effect was also reported to be mediated by collagen VI-dependent changes in mechanical properties, which are not different in our ECM hydrogels. Thus, it is difficult to deconvolute the contributions due to mechanical versus biochemical differences. Glycosaminoglycan and proteoglycan composition is also known to be important for the satellite cell niche due to their ability to bind and sequester growth factors [50-52]. Thus, differences in proteoglycan composition between the two materials could also have an effect on regeneration *in vivo.* Previous studies have shown that the skeletal muscle ECM hydrogel may act directly on skeletal myoblasts by increasing proliferation and differentiation [8, 40], so a complex distribution of proteins and proteoglycans may be more important than the presence or absence of a single component. Studies have shown that decellularized ECM hydrogels are more regenerative than a single ECM component such as collagen [13, 28], which supports this finding *in vivo*.

ECM hydrogels have been shown to alter the inflammation and metabolism response in endogenous tissue [8, 53]. Polarization of the inflammatory response can have a large effect on regeneration *in vivo* [31, 54]. It is unclear which ECM components specifically can cause varying inflammatory responses but observed differences between the endogenous response to skeletal muscle and lung ECM hydrogels show this does occur. It is possible that some of these effects may be caused by artifacts of the decellularization process. In this experiment, we controlled for decellularization detergent, which is known to effect the final ECM composition [55], to avoid overt differential influences, however, this still could have been a factor as it is practically impossible to utilize the exact same protocol given tissue differences. For example, the skeletal muscle ECM required a higher SDS concentration than the lung for effective decellularization.

This study was the first to show that endogenous regeneration response in the context of ECM hydrogel injection is not interchangeable with respect to source of ECM injected. Distinct upregulation of muscle contractility genes, along with supporting histological evidence of skeletal muscle progenitor cell recruitment, activation, and shift in increased fiber area all indicate that tissue specific ECM hydrogels are critical for inducing tissue specific regenerative responses *in vivo*, at least in the case of skeletal muscle. Optimal regeneration with tissue specific ECM hydrogels may depend on application and design goals, and thus ECM choice is an important consideration for tissue engineering approaches.

## 5. Conclusions

In this study, we demonstrated that a skeletal muscle ECM hydrogel induced beneficial effects on muscle regeneration and hypertrophy after acute injury *in vivo.* This data indicates a potential role for skeletal muscle-specific regenerative capacity of decellularized, injectable skeletal muscle hydrogels. Because viscoelastic properties of the hydrogels were similar, it is likely that biochemical differences between the materials guided the differences in endogenous cellular behavior and increased muscle regeneration *in vivo.* While specific clinical applications may differ, this study suggests ECM choice is an important consideration for optimal tissue regeneration.

## Data availability

All relevant data is available upon request from corresponding author.

## Acknowledgments

We would like to thank Kristen Jepsen and the Institute for Genomic Medicine at UCSD for their help with library prep and RNA sequencing. This work was supported by the California Institute for Regenerative Medicine (TRAN1-09814) and the National Institutes of Health (NIH) Heart, Lung, and Blood Institute (NHLBI) (R01HL113468). J.L.U. was supported by a NIH NHLBI F31 fellowship (F31HL136082).

